# High thermal plasticity and vulnerability in extreme environments at the warm distributional edge: the case of a tidepool shrimp

**DOI:** 10.1101/2020.04.25.061424

**Authors:** Eyal Amsalem, Gil Rilov

## Abstract

Climate change threatens the resilience of species, especially at their warm distributional edge in extreme environments. However, not much is known about the thermal vulnerability of marine intertidal species at this edge. We investigated the thermal vulnerability of the tidepool shrimp, *Palaemon elegans* in the fast-warming southeastern Mediterranean, its warm distributional edge. Tidepool organisms experience strong and fast thermal fluctuations. This might make them more resilient to change, but also bring them closer to their thermal limits during extreme conditions. To test the shrimp’s resilience, we tested three hypotheses: (1) *P. elegance* in the southeast Mediterranean has higher critical thermal maximum (CTMax) than in cooler regions, (2) the shrimp possess seasonal acclimatization, but (3) long exposure to extreme summer temperatures might erode its thermal performance making it vulnerable to future climate change. We characterized the shrimp’s thermal environment and population dynamics, determined CTMax and tested diverse physiological performance attributes (respiration, digestion, activity, growth) under a wide range of temperatures during winter and summer. *P. elegans* has a wide optimum performance range between 20-30*°*C during summer and its CTMax is 38.1°C, higher than its Atlantic counterparts. However, its warming tolerance is only 0.3°C, indicating low capacity for dealing with further warming in pools compared to northeast Atlantic populations that have wider tolerance. Prolonged exposure to current mean summer values in open water (∼ 32*°*C) would also significantly reduce its performance and increase mortality. This suggests that its population viability may be reduced under continuous regional warming and intensification of extreme events.

## 2. Introduction

There is a growing body of evidence concerning the increasing impacts of anthropogenic climate change (CC) on marine ecosystems and their biodiversity, which is predicted to continue and increase in the near future (Gattuso et al., 2015). Ocean warming poses one of the biggest threats to sea life, with evidence that the ocean surface has already warmed by a global average of 0.87°C compared to the pre-industrial period, and is expected to increase further by 1-3°C, depending on the emission scenario (IPCC, 2019). Temperature affects all biological and chemical processes and ectotherms are directly affected by temperatures since they lack the ability to control their own body temperature. Effects of temperature change at the organismal level, i.e., physiology and behavior (which affect growth, reproduction and levels of activity), ultimately lead to impacts on species global distribution, seasonal timing of biological events (phenology), and food web structure (Pörtner and Farrell, 2008; Verberk et al., 2016). At the physical-chemical-physiological interface, increasing temperature leads to increases in the kinetic energy of molecules, and therefore increasing metabolic rates such as enzymatic reaction as well as mitochondrial activity that in turn intensifies the respiration rate of an organism (Levinton, 2001; Schulte et al., 2011).

Temperature increase under global warming is heterogeneous in space and time and some regions warm faster than others (Macias et al., 2013). Because in many locations or marine habitats organisms already live close to their thermal limits (Harley et al., 2006; Somero, 2002; Vinagre et al., 2018) even slight increases in temperature can negatively affect their performance, and ultimately their survival (Pörtner and Farrell, 2008; Rivetti et al., 2014). The warm or trailing edge of species distributions is where species are expected to be frequently thermally challenged, and where the viability of the population should start to erode first under global warming (Bates et al., 2014; Donelson et al., 2019; Somero, 2010). At the warm edge, even thermal refugia can be lost in extreme environments such as the intertidal zone (Lima et al., 2016).

The southeastern Mediterranean Sea, represents the warm edge of most Atlanto-Mediterranean species and can be used as a testbed of marine species vulnerability at this edge. The Mediterranean Sea, as a whole, experiences one of the strongest ocean warming rates because of its small size and the fact that it is almost enclosed, in addition to its geographic location and the effect of a recent change in the Atlantic Multidecadal Oscillation (Macias et al., 2013; Marbà et al., 2015). Therefore, the Mediterranean is expected to be more strongly influenced by global change than the open ocean, and indeed there is accumulating evidence about the footprint of climate change on Mediterranean Sea biota, mostly for its well-studied western basin (Marbà et al., 2015; Rivetti et al., 2014). For example, the two major mass mortality events that followed heat waves in the northwest basin in the past two decades (Rivetti et al., 2014). The East Mediterranean Sea (EMS) is warming faster than the western basin, especially in winter (Pastor et al., 2019), and offshore *in-situ* measurements over the past three decades revealed warming at an average rate of 0.12°C per year or almost three degrees in the past three decades (Ozer et al., 2016), an order of magnitude larger than the global average. Nonetheless, the near absence of long-term biological time-series makes it difficult to identify the ecological footprint of climate change in this basin. Recently, the collapse of many native reef invertebrate species has been reported on the southeastern coast of the Levant basin (Israel), and it was suggested that rapid ocean warming in this region may be partially responsible for that (Rilov, 2016). At least for sea urchins, which have become extremely rare on this coast in the past few decades, there is evidence that they die when water temperature rises above 30.5°C, which occurs every summer on this coast in the past decade (Rilov, 2016; Yeruham et al., 2015).

One of the ecosystems where climate warming is expected to have a strong influence is the rocky intertidal zone. This zone exists at the margin between the terrestrial and marine realms, so organisms in it are subjected to frequent changes in both water and air temperatures. Within this ecosystem, tidepools may be especially vulnerable to change in temperatures (Harley et al., 2006; Helmuth et al., 2006). Tidepools are highly structured environments that provide food and shelter for diverse biological communities (Vinagre et al., 2015). Each tidepool has its own unique environmental characteristics, since they can be found in a variety of shore levels, sizes and volumes, as well as having different levels of connection to the open sea (Metaxas & Scheibling 1993, Vinagre et al., 2015). Because of their small water volume, tidepools have low thermal inertia, unlike other aquatic environments, and can thus warm or cool very fast, reaching extreme values in a matter of hours (Vinagre et al., 2018). This requires organisms inhabiting them to be able to deal with a large range of abiotic conditions, but also possibly make them more prone to extreme change in their fitness (Madeira et al., 2012b). A limited global analysis across realms suggested that animals from more stable environments have greater capacity for acclimation compared to unstable ones, but pointed out that generality of such patterns is limited because much of the world is undersampled (Seebacher et al., 2015).

On the rapidly warming Israeli coast, which is the southeastern edge of distribution of most Mediterranean and Atlanto-Mediterranean species, and where the water is naturally hottest in the basin, tidepool ecology has not been studied before, but it is assumed that the pools can get extremely hot when disconnected from the sea. This study focuses on the small rockpool shrimp *Palaemon elegans* (Rathke 1837). *P. elegans* is native to the Mediterranean Sea, the Black Sea, and the East Atlantic coast, ranging from Norway in the north to Namibia in the south. It inhabits coastal tidepools, shallow rocky areas, submerged vegetated areas, lagoons, estuaries and man-made structures (Bilgin et al., 2009; Glamuzina et al., 2014). Its reproductive season is May to September (in the Black Sea), and when conditions are favorable, the female can reproduce twice a year (Zaharia et al., 2005). The shrimp is euryhaline, with a high salinity tolerance of 1-40 PSU (Glamuzina et al., 2014). It is omnivorous, feeding predominantly on algae, small crustaceans and foraminifera (Dalla Via, 1985; Glamuzina et al., 2014). In Israel, it almost exclusively lives in tidepools on the rocky shore (Tsurnamal, 1963). *P. elegan*s distribution expanded extensively in the past century, as it had invaded the Caspian Sea (1930), Aral Sea (1960), Persian Gulf (1972), Baltic Sea (2000) and the shores of Massachusetts, North America (2010) (Fofonoff et al., 2003). As a result, the species gained widespread attention in the fields of ecology, biology, and in relation to commercially-related impacts on food security and its use as bait in fisheries (Al-Khafaji et al., 2016; Janas and Mańkucka, 2010; Janas et al., 2013). Other studies from the Black Sea, Caspian Sea and the north Mediterranean Sea investigated its growth and reproductive performance, larval development, fecundity, individual and group behavior, population structure and habitat conditions (Bilgin et al., 2009; Dalla Via, 1985; Duran et al., 2006; Glamuzina et al., 2014; Janas and Mankucka, 2010; Tsurnamal, 1963). A few studies showed that *P. elegans* has a strong resistance and a high tolerance to a wide range of conditions. It achieves this by exhibiting a high production of heat shock proteins (HSP70) and by expanding its habitat range through occupying vacant ecological niches (Madeira et al., 2012a; Szaniawska and Lapinska, 2006; Tsurnamal, 1963). Other studies examined the vulnerability of this species (among other species) to global warming in the Northeast Atlantic and in the west Mediterranean Sea, by investigating the CTMax and its acclimation capacity through different acclimation states at different life stages (Madeira et al., 2015; Madeira et al., 2012b; Magozzi and Calosi, 2015; Ravaux et al., 2016; Vinagre et al., 2013; Vinagre et al., 2016) (Table 1).

**Table 1.** CTMax values of *Palaemon elegans* tested in studies conducted in the north east Atlantic and west Mediterranean Sea and determined under different temperature exposure regimes. Acl = Acclimation tretment, D = days.

| Authors | Location of collection | Time of collection (mm-yy) | Water temp at collection time | Maximum habitat temp | Heating rate | *P. elegans* CTMax |
| --- | --- | --- | --- | --- | --- | --- |
| Diana et al | Portugal | Jul-10 | 24°C | 35°C (based on max air temp) | 1°C\H | 34°C |
| Magozzi & Calosi | England | Jun-12 | 12.4°C | Not reported | 0.75°C\ M | Acl 10°C: 30°C<br>Acl 15°C: 33°C<br>Acl 20°C: 34°C<br>Acl 25°C: 35°C |
| Catarina et al | Brazil and Portugal | Sep-14 | 20°C | 30°C (June 2014) | 1°C\H | 7D: 33.4°C<br>30D: 36.2°C<br>10D: 36.6°C |
| Juliette et al | France | Oct-11 | 17°C | Not reported | 1°C\M | Acl 10°C: 27.5°C<br>Acl 20°C: 33.2°C |

Here, we investigate the effects of temperature on *P. elegans* at its distributional trailing (warm) edge, the EMS, where it possibly experiences the most extreme conditions in both open water and tidepools. A species’ relative sensitivity to warming depends on its thermal tolerance and also on its thermal acclimation potential, defined as the plasticity of its behavioral, physiological or morphological characteristics in response to environmental temperature (Angilletta Jr, 2006). Hence, the present study specifically aims to investigate the vulnerability of the EMS population of *P. elegans* to the predicted rise in temperature. For that, we measured temperature and population dynamics in different pool types occupied by the shrimp, and tested its CTMax and thermal performance of various physiological and behavioral parameters under a wide range of temperatures and at different seasons (for possible seasonal adaptations). Information regarding the shrimp’s optimal thermal performance range and its upper thermal limits (CTMax) will help to evaluate the present and future effects of CC on the species’ fitness and ultimately its distributional range. The thermal performance curve (TPC) describes the effects of temperature on the rate of biological processes, and demonstrates the thermal range of performance, the optimal temperature (T_opt)_ and the points at which performance is zero (T_crit_). CTMax explores the upper thermal tolerance of a species and is a good index and standard for evaluating the thermal requirements and physiology of an organism (Lutterschmidt and Hutchison, 1997). We specifically tested three working hypotheses: (1) CTMax of the southeast Mediterranean shrimp is higher than that of the East Atlantic one because of the extreme environmental condition it experiences in the EMS (Vinagre et al., 2016); (2) T_opt_ will be higher in the summer than in the winter as a result of seasonal acclimatization, which makes the shrimp more tolerant to higher temperatures in the warm months (Chapperon *et al*., 2016), and (3) longer exposure to high temperatures will erode the performance of the shrimp, reducing its optimum values.

## 3. Materials and methods

### 3.1 Study site

We studied the shrimp population at the beach of the Shikmona marine reserve, Haifa, Israel (Fig. 1, 32° 49’ 32.27’N, 34° 57’ 18.21’E). This site has a gently-sloping limestone rocky shore at the back of a vermetid reef platform, with multiple tidepools (rockpools) of varying sizes and connectivity to the open waters. The environmental conditions and population dynamics of *Palaemon elegans* were studied in five tidepools of different sizes, and varying degrees of connection to the open sea (Fig. 1).

**Figure 1.**
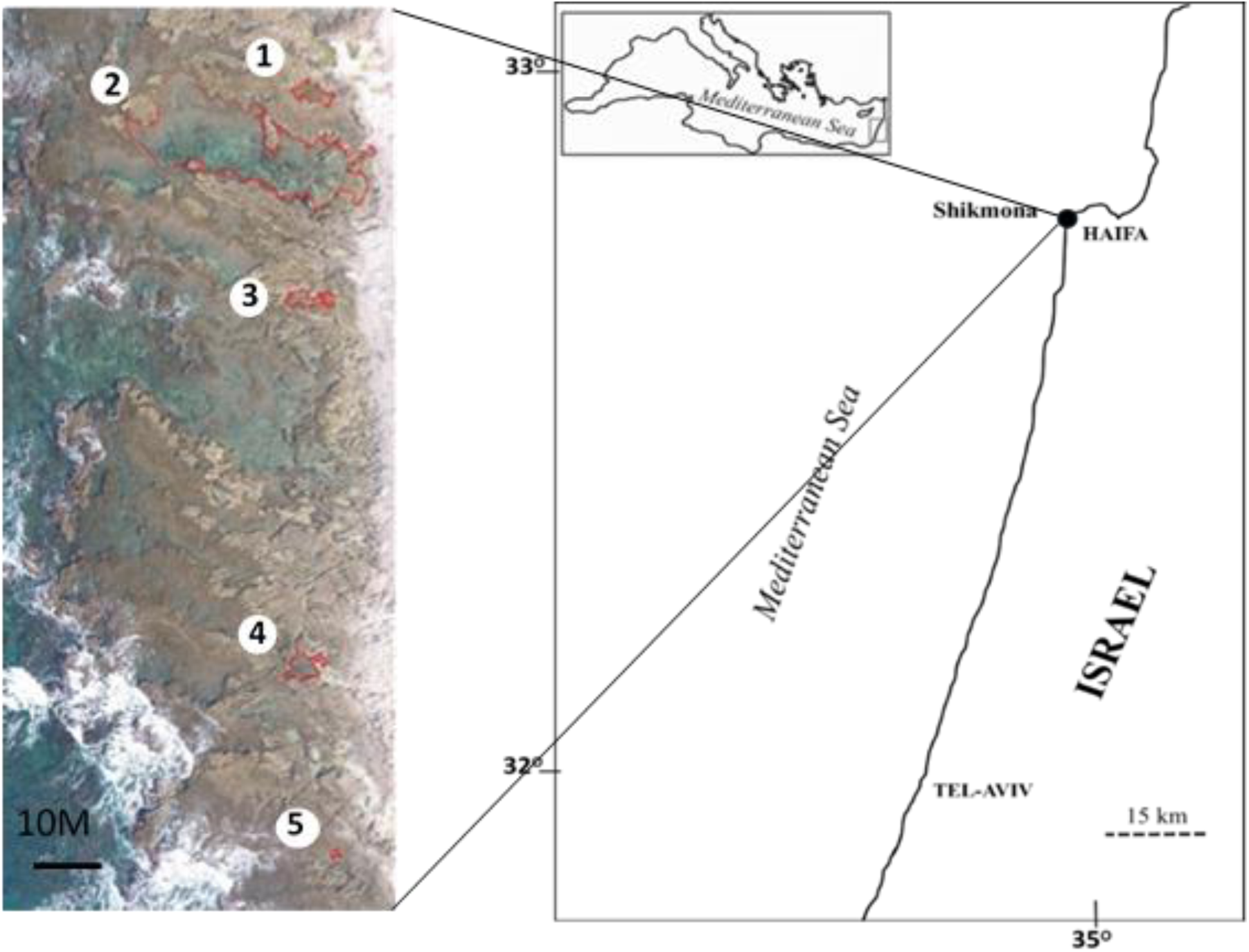
Study site. Location of the study site on the Israeli coast and a drone photo showing the location of the five monitored tidal pools on the rocky shore at Shikmona.

### 3.2 Environmental conditions

In order to determine the dynamics and annual range of temperatures the study species is naturally exposed to in tidepools in the study region, water temperatures were recorded hourly between November 2016 and January 2018, using HOBO pendant temperature Data Logger (model UA-002-64, Onset Computer Corporation, USA) deployed at the bottom of the five pools and attached to the rock with stainless-steel bolts similar to Kelly et al (2012). Measurements of nearshore open-water temperatures were obtained from a logger deployed at a depth of 0.5 m at the seaward edge of the rocky shore (serving IOLR’s National Monitoring Program). Measurements of air temperatures were obtained from a weather station located on the roof of IOLR, about 200 m from the study site.

### 3.3 Population dynamics

In order to detect seasonal population dynamics and determine the reproductive and recruitment season, the population of *P. elegans* was sampled monthly in the five tidepools between December 2016 and December 2017 using a hand net (12 × 10 cm, 1mm mesh). The net was moved slowly along the sides (walls) and near the bottom of the pool (areas where the shrimp can normally be found) for three minutes, and all individuals caught were recorded and placed back to the same pool after handling. Because this species is nocturnal, the sampling occurred at night when the sea is calm and during low tide. The recording included measuring the size with a digital caliper (INSIZE, electronic caliper series 2111) and weight with an analytical scale (Analytical balances ASB-220-C2 (±0.2mg)).

### 3.4 Specimen collection and preparation for experiments

Specimens of *P. elegans* were collected from tidepools (different pools than the monitored ones, but at the same site, from depths of 0.1-0.5m) for the different experiments, using a hand net in January, May, June and September of 2017. The average water temperature in the tidepools at the time of collection was 17°C, 22°C, 25°C and 29°C, respectively. Immediately after collection, the shrimp were transferred to the lab for an initial acclimation period of five days in outdoor aquaria (20×38×21cm, maximum 25 shrimp in each aquarium) with a continuous seawater flow at ambient conditions (supply directly from the sea).

### 3.5 Temperature performance experiments

Different measurements of the shrimp’s thermal performance under a range of temperatures were taken three times: adults were tested in winter and summer and juveniles during the main reproductive season in spring. After the acclimation period in the aquaria, the shrimp were transferred to individual glass jars set in ten separate 20L units of an outdoor (allowing natural photoperiodicity) custom-made thermobath system (see description in Guy-Haim et al., 2016) for an additional five days of acclimation at a constant ambient temperature, corresponding to the average seawater temperature measured at the sampling site at each season. The temperature in each thermobath was controlled independently by NOVUS N1020 temperature controllers (±0.4°C). Individuals of *P. elegans* were each placed in separate 0.75L glass jars (n=6) that were submerged in the thermobaths. Every 48 hours, 100% the water was replaced manually and gradually in each container with filtered Mediterranean seawater (2µm carbon filter) that was pumped from a depth of 1m and warmed or cooled beforehand to the temperature currently present in each thermobath treatment in order to reduce possible stress. Food supply was *ad libitum*. Before placement in the jars, each shrimp was weighed with an analytical scale (Analytical balances ASB-220-C2 (±0.2mg)) after being carefully dried with blotting paper. Salinity was kept at the levels of the nearby coastal water and was on average 39.0±0.1 PSU. Water motion and temperature homogeneity within the containers were ensured by aerators and submersible pumps. After five days of the acclimation period, the temperature in each bath was differentially increased or decreased by 0.5°C·12h^-1^ until reaching their target temperatures at the same date. This rate of change was not greater than the diurnal temperature range in the collection site at that period (1°C). The winter experiment (temperature range: 8-34°C) started on February 4^th^, 2017; target temperatures were reached on February 24^th^, 2017 (25 days after sample collection); and were maintained until the end of the experiment, on March 17^th^, 2017. The late spring juvenile growth experiment (temperature range: 12-35°C) started on June 2^nd^, 2017; target temperatures were reached on June 20^th^, 2017 (23 days after sample collection) and were maintained until the end of the experiment, on June 26^th^, 2017. The late summer experiment (temperature range: 17-35°C) started on September 25^th^, 2017; target temperature was reached on October 10^th^, 2017 (20 days after sample collection) and was maintained until the end of the experiment, on October 21^st^, 2017. We tested the performance of four types of animal functions that could be affected by temperature, and thereby can affect the shrimp’s fitness: respiration, digestion rate, growth (in juveniles) and activity (movement).

#### 3.5.1 Respiration

was measured as the change in dissolved oxygen (DO) during 1-3-hour incubations. The incubations were performed three times on individual specimens in gas-tight 0.3L containers to which the shrimp were temporarily relocated. DO concentration (µmol O_2_ ·L^-1^) was measured inside the containers using an oxygen optode (Oxi3315, WTW Weilheim, Germany). The first incubation was done after a five-day acclimation period at ambient temperatures, the second, four days after reaching the target temperatures and the last - a week after the second incubation, in order to see if longer exposure had shifted the shape of the performance curve as suggested by our third hypothesis. In the winter experiment, a fourth incubation was done, ten days after the third incubation. The shrimp were weighed after each incubation. Net respiration in terms of oxygen consumption (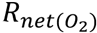 in units of µmol O_2_ ·liter seawater (L) ·gram shrimp (GS) ·hour (H)^-1^) was calculated as the difference between initial and final dissolved oxygen concentrations (*ΔDO*) per liter seawater per gram of shrimp divided by the duration of the incubation (*ΔT*) as follows:

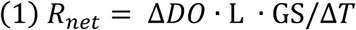

#### 3.5.2 Gut resident time

(GRT) as a measure of digestion rate was tested three days after the second incubation. Food in the shrimp’s gut is easy to follow because of its highly transparent body. Before the test, the shrimp were starved for two days and their glass container was cleaned every day from shrimp feces and small algae that grew on the jar sidewall. After this period, the digestive organs of the shrimp were clearly empty. Then, each individual received five red pellets of fish food (SERA Granured), and all shrimp were continuously observed to detect the first time they defecated. GRT = the interval between feeding and first defecation.

#### 3.5.3 The growth rate

of juveniles (smaller than 7mm) was assessed by measuring their weight and size in the beginning of the experiment and once every two weeks for two months. This was done by weighing the shrimp with an analytical scale (Analytical balances ASB-220-C2 (±0.2mg)), and photographing them with a digital camera (Sony-RX100, 5) to assess size change with the Image-J software.

#### 3.5.4 Movement rates (activity)

were measured using a method similar to the one described by Chapman et al (2013) with small modifications. We used a 19×24×17 cm aquarium that was divided into 12 equally-sized square grids (6.3×6 cm). Individual shrimps were placed in the aquarium for three minutes of acclimation, then their movement was recorded with a digital video camera (Sony-RX100, 5) for another three minutes. We considered movement as follows: every time the shrimp moved from one square to the next or moved upward. The activity rate was defined numerically as the number of movements.

### 3.6 Critical temperature maximum (CTMax) and vulnerability assessment

The thermal tolerance of the shrimp was determined by the dynamic method described by Mora and Ospina (2001). The aim was to determine the CTMax, which is defined as the “arithmetic mean of the collective thermal points at which the end-point is reached” (Mora and Ospina, 2001). The end-point was a loss of equilibrium (LOE) when the shrimp starts to swim upside-down. To determine the CTMax, individual shrimp were placed in an aquarium (19×20×38 cm) with a water depth of 13 cm. Water was warmed at a constant rate of 1°C h^-1^ using an ENDA EPC8420 profile controller with SCHEGO Heizer 300-watt heater and a temperature sensor. The shrimp were observed continuously until they reached the end-point. The end-point temperature was recorded and then the CTMax was calculated as follows:

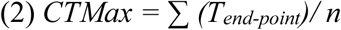

Where T_end-point_ is the end-point temperature for individual 1, individual 2, individual n, divided by n individuals that were tested (n=X). The warming tolerance was determined as the difference between CTMax and the maximum habitat temperature (MHT).

### 3.7 Survival

While *P. elegans* were in the thermobath system, their survival was measured from the point when they got to the target temperature. The daily survival rate of *P. elegans* in the experiment was calculated as follows:

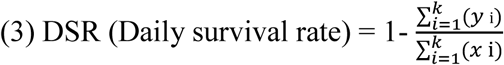

Where K is the total number of animals in the sample, *yi* is 1 if animal *i* died during the reporting period and 0 if animal *i* was still alive at the end of the reporting period, *xi* is the number animal days (number of day the individual was alive) for animal *i* (DeMaster and Drevenak, 1988).

### 3.8 Establishing performance curves

The performance parameters were plotted against temperature and fitted with models commonly used to describe performance curves (Angilletta Jr, 2006). The corrected Akaike Information Criterion (AICc), estimating information loss upon model selection, was calculated for each model, where the lowest AICc scores indicate the best fitting model. AICc and r^2^ indicators were used to assess the goodness-of-fit of thermal performance curves. Data exploration was performed to find P_max_ (the value of the measured parameter at peak performance) and TOPT (the optimal temperature where performance levels are maximal). The assessment of model fitting was performed using CurveExpert Professional 1.6.10.

### 3.9 Statistical analysis

One-way analysis of covariance (ANCOVA) was performed to test the effect of treatment-temperature and season-winter/summer and their interaction with respiration, GRT and movement. For the effect of time since reaching the target temperature on respiration rate, one-way repeated measures ANOVA was conducted. For growth rate, differences between treatments were tested using one-way ANOVA. A subsequent Bonferroni post-hoc test was applied to reveal which treatment differs from the others, and how. All assumptions were tested prior to the analysis and a significance level of 0.05 was used. In cases where samples were not normally distributed (Shapiro Wilk’s test) or not homoscedasticity (Levene’s test), data was first transformed using Log(10) transformation, and in cases where prerequisites for parametric tests were not achieved, the non-parametric Kruskal-Walis test was performed. Statistical analysis did not include dead shrimp. All statistical tests were performed using SPSS statistics version 21. Tabular results of the statistical analyses can be found in the Supplementary Material section.

## 4. Results

### 4.1 Tidepools and open sea temperatures

Pool depth varied between 20 and 95 cm and their area ranged from 0.72 to 367.3 m^2^ (Supplementary Table S2). The temperature varied considerably both daily and seasonally between pools (Fig. 2. A-F), with maximum daily fluctuations in the closed pools reaching 10°C. At summer time, in closed disconnected pools (e.g. pool 1) the temperature raised above 35°C, for short periods of 6-7 hours maximum and cooled at night (Fig. 3). Minimum and maximum extreme values of 5.9°C in November and 37.8°C, respectively, in July were measured at pool 1. In the shallow open coastal waters near the reef, the minimum temperature was 14.9°C in December and the maximum was 32.18°C in July (Fig. 2G) and maximum daily fluctuations were 3°C during the sampled year.

**Figure 2.**
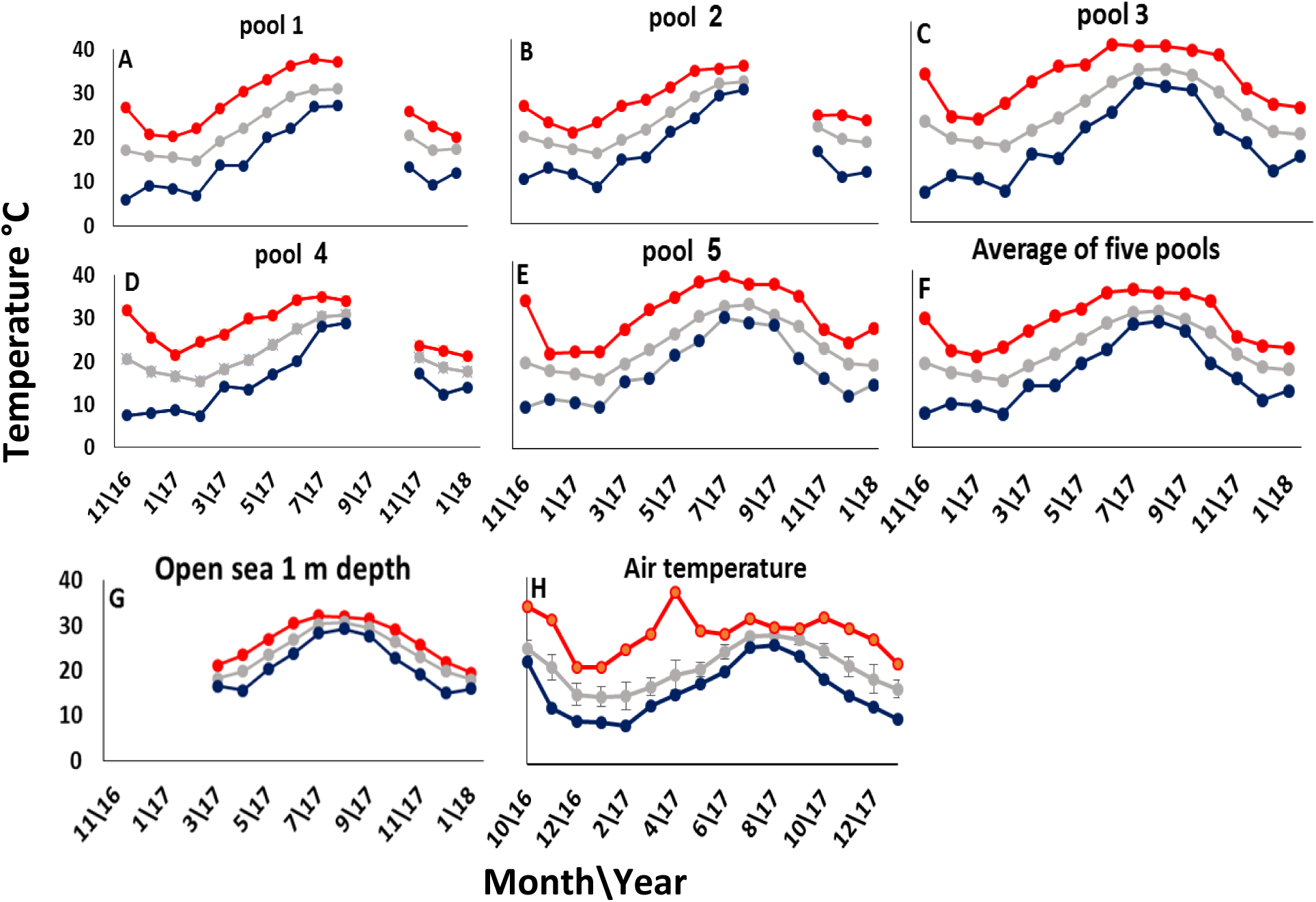
Tidepool monthly temperatue variability. (**A**)-(**E**) Mean monthly water temperature in the five pools during the study period. (**F**) Average temperature of five pools in the study site. (**G**) Coastal temperature in the open water at 0.5 m depth during the study period. (**H**) Air temperature during the study period. Red, gray and blue lines represent the maximum, average and minimum values that measured, respectively (gaps in data are periods when the logger failed and replaced later). In three pools, loggers were lost or damaged resulting in gaps in data.

**Figure 3.**
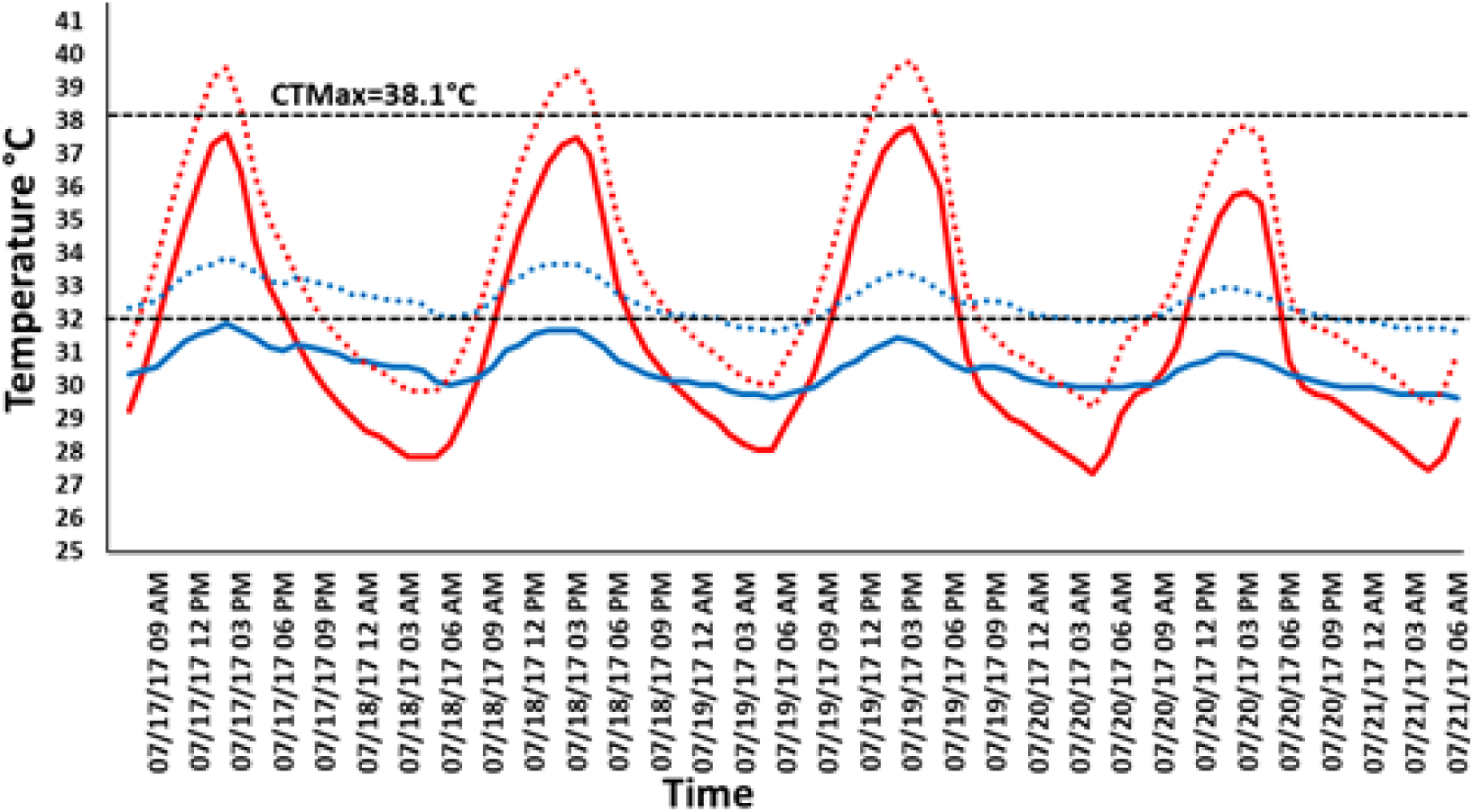
Hourly pool temperature variability. Four days of temperature fluctuations measured in July 2017 illustrating the present and predicted future summer conditions in the most extreme (isolated tidal pool 1, solid red line) and benign (open sea, solid blue line) habitats. The red and blue dashed lines represent increase of +2°C in water temperature. The upper black dashed line represents the measured critical temperature maximum and the lower line represents the measured threshold point for *P. elegans* after four days of exposure to constant temperature (32°C).

### 4.2 Population dynamics

Individuals of *Palaemon elegans* were observed in the monitored tidepools throughout the year. The highest number of *P. elegans* was caught in tidepool 1 (remote and disconnected) and the lowest in tidepool 2 (big open pool). Ovigerous females were present from March-April and July-October. Juveniles were present between May-June and September-October. The highest number of juveniles were caught in May in tidepool 1 (Supplementary Table S1).

### 4.3 Thermal performances

#### 4.3.1 Respiration

In the first incubation (at ambient water temperature) of adults in each season, we examined the relationship between the respiration rate (µmol O_2_ h^-1^) and the shrimp biomass (in grams). In both seasons, the expected increase in the respiration rate with body mass (see, Ivleva 1980) was evident (winter: r^2^ = 0.45, summer: r^2^ = 0.64, Supplementary Fig. S1). The best fitting models for the performance parameters measured after acclimation to target temperatures are detailed in Table 2.

**Table 2.** Best-fitting models for the tested TPCs in different seasons and for the performance of respiration at different exposure durations after reaching target temperatures. The goodness-of-fit indicators are Akaike Information Criterion (AICc) and r^2^.

| Performance variable | N | Best-fit model | AICc/$r^2$ | T<sub>OPT</sub> (°C) | P<sub>max</sub> | CT<sub>Max</sub> |
| --- | --- | --- | --- | --- | --- | --- |
| Respiration, 4 days, winter | 58 | Rational | 131.6/0.6 | 28.2 | 11.3 | 33.5 |
| Respiration, 11 days, winter | 58 | Rational | 122.7/0.56 | 25.5 | 9.5 | 33.5 |
| Respiration, 20 days, winter | 53 | Rational | 51.86/0.45 | 25.5 | 4.8 | 34.5 |
| Respiration, 4 days, summer | 49 | Rational | 166.4/0.35 | 32 | 14.4 | 35.5 |
| Respiration, 11 days, summer | 49 | Rational | 152.66/0.54 | 26.3 | 14.7 | 34.8 |
| Activity winter | 42 | Rational | 261.4/0.08 | 29 | 33 | 34.5 |
| Activity summer | 42 | Heat capacity | 260.09/0.1 | 26-30 | 40 | N/A |
| Growth | 64 | Rational | 1141.3/0.3 | 24 | 0.0003 | N/A |

When dead individuals are included in the analysis we see a clear optimum that is shifting with the duration of exposure and with season (Fig. 4). When dead individuals are excluded, a steady increase in respiration rate with temperature is evident, with some leveling or even slight decrease (Fig. 4D-E) at the higher temperatures (insets in Fig. 3). A clear reduction in T_opt_ of several degrees, with the increase in duration of exposure to target temperature, was evident in both winter and summer (Table 2, Fig. 4). In winter, P_max_ was also reduced considerably with duration of exposure (Table 2, Fig. 4). Significant differences were found in respiration rate between winter and summer, after four days in the target temperature (incubation 2), with an interaction between season and temperature (ANCOVA, P < 0.05, Supplementary Table S3). Pairwise-comparisons for the winter incubation 2 (Kruskal-Wallis, Supplementary Table S4) show significant differences (P < 0.05) between 8°C and 25°C, 28°C, 30°C and 32°C. Pairwise-comparisons for summer incubation 2 (Bonferroni post-hoc test, Supplementary Table S5) show significant differences (P < 0.05) between 17°C and 28°C, 30°C, 31°C, 32°C, 33°C, 34°C, and between 21°C and 31°C, 32°C and 33°C. The same result (interaction effect of season and temperature) was found after eleven days in the target temperature (ANCOVA, P < 0.001, Supplementary Table S6). Pairwise-comparisons for winter incubation 3 (Bonferroni post-hoc test, Supplementary Table S7) show significant differences (P < 0.05) between 25°C, 8°C and 12°C. Pairwise-comparisons of temperature for summer incubation 3 (Bonferroni post-hoc test) did not show significant differences between temperatures.

**Figure 4.**
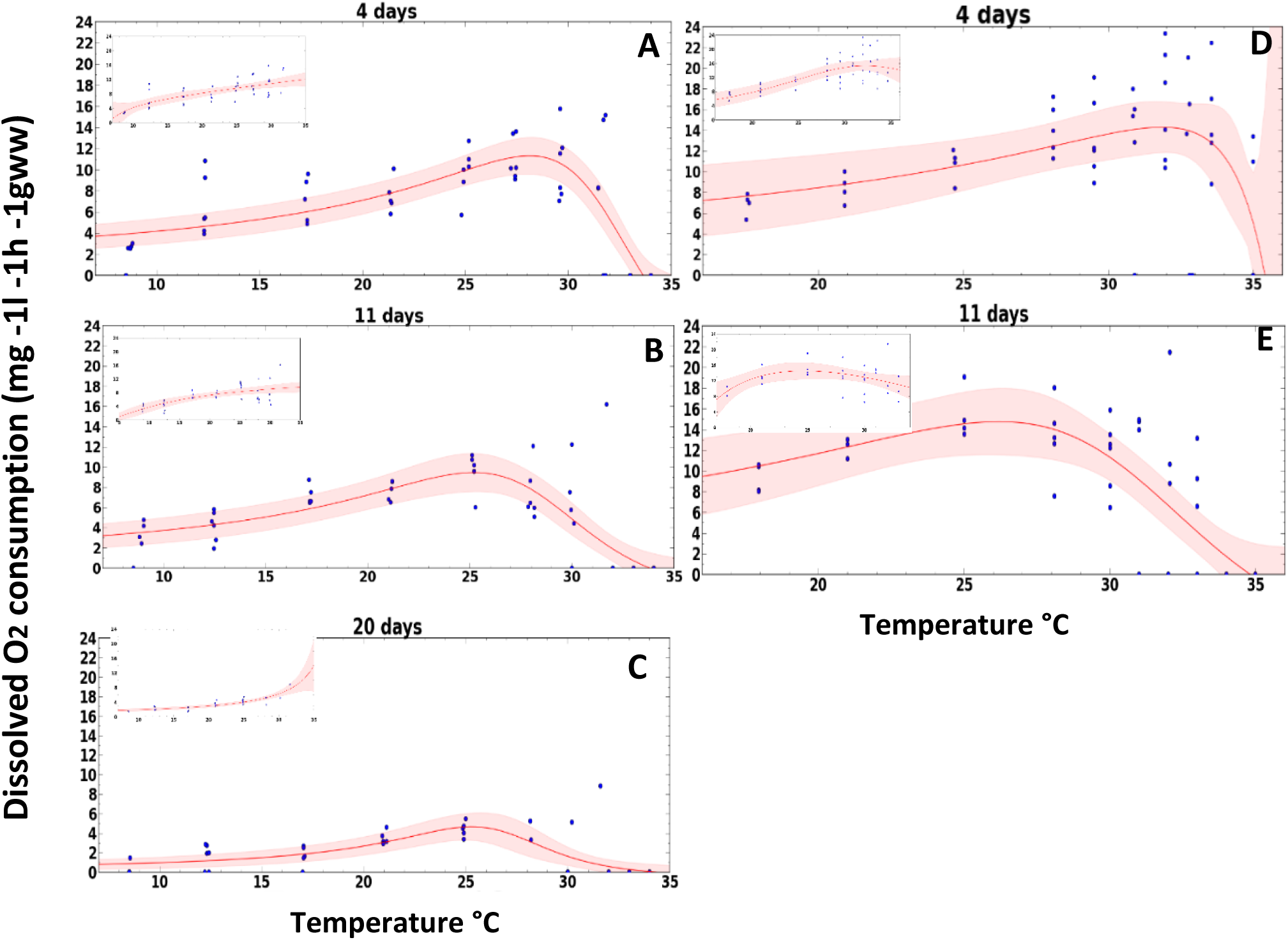
The effect of temperature on respiration. Temperature effects on the dissolved oxygen (DO) consumption in *P. elegans* measured at 10 temperature levels in the winter between 8-34°C and in the summer between 17-35°C (n=6 per temperature level). Measurements were conducted after reaching the target temperature. In the winter: after 4 days (**A**), after 11 days (**B**) and after 20 days (**C**). In the summer: after 4 days (**D**) and after 11 days (**E**). A 95% confidence interval bend is marked by shaded areas. Insets show the data and curves when all dead individual are removed. The model best fitting to the data were rational (**A-D**) and Heat capacity (**E**).

The effect of duration of exposure in the winter experiment (Repeated measured, P < 0.001) was significant (Bonferroni post-hoc test, P < 0.05) (Supplementary Table S8) between 12°C and 21°C, 25°C and 28°C, as well as between 17°C and 25°C. The effect in the summer had significant differences (P < 0.05) between 17°C and 31°C and 32°C (Supplementary Table S9).

#### 4.3.2 Gut residence time (GRT)

In both seasons, there was a strong negative correlation (r^2^=0.644) between temperature and GRT (Fig. 5), but at the highest temperature treatments (winter: 34°C, summer: 35°C) shrimp showed low interest in food or did not eat at all (thus data are not shown for these temperatures). GRT in the summer was much lower in all tested temperatures compared to winter (Fig. 4a). There was a significant difference between the interaction of season and temperature (ANCOVA, P < 0.001, Supplementary Table S10). Pairwise-comparisons of temperatures were done separately for winter and summer and showed significant differences (Kruskal-Wallis, P < 0.05, Supplementary Tables S11, S12): in winter between 28°C and 8°C and 12°C, in summer: between 25°C and 33°C.

**Figure 5.**
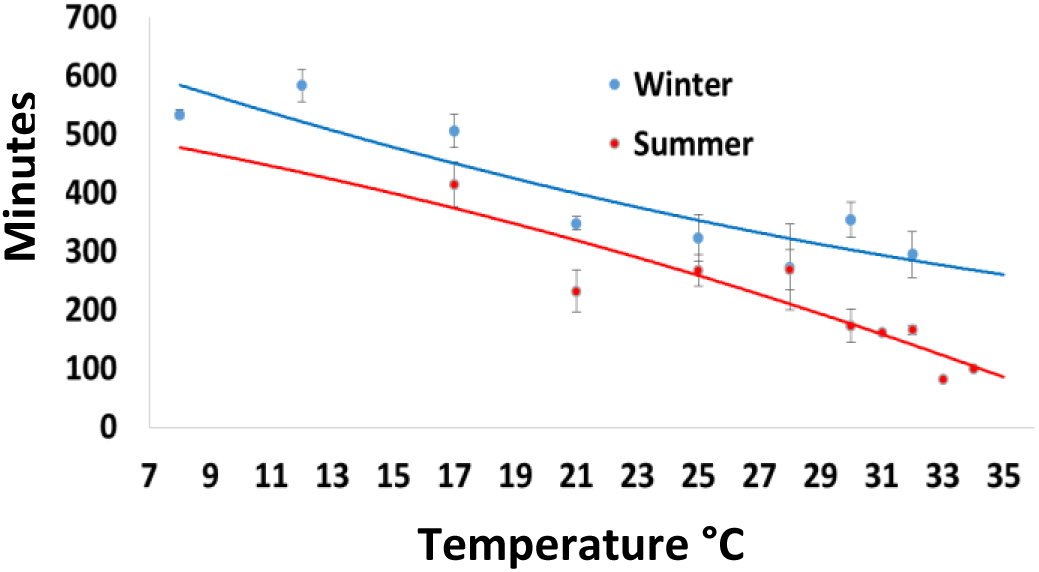
The effect of temperature on the gut residence time in *P. elegans* measured in the winter and summer at 10 temperature levels between 8-34°C and 17-35°C, respectively (n=6 per temperature level). Resident time is shown as means ± SE.

#### 4.3.3 Shrimp activity

Activity was highly variable among individuals at all temperatures, resulting in a low power of analysis (Fig. 6). Nonetheless, the interaction between season and temperature was significant (ANCOVA, P < 0.05, Supplementary Table S13), but pairwise-comparisons, done separately for winter and summer, found significant differences only in the summer, between 17°C and 33°C (Bonferroni post-hoc test P < 0.05, Supplementary Table S14).

**Figure 6.**
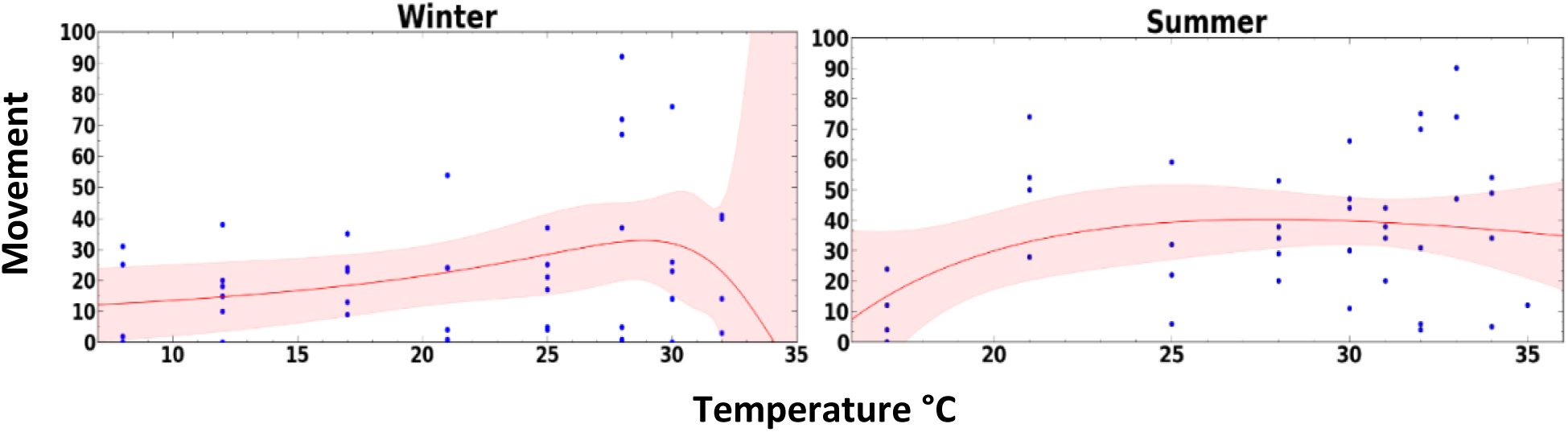
The effect of temperature on activity of *P. elegans* measured in the winter and summer at 10 temperature levels between 8-34°C and 17-35°C, respectively (n=6 per temperature level). Acitvity mmeasurements that were conducted six days after reaching the target temperature. Each dot represents a measurement. A 95% confidence interval bend is marked by shaded areas. The fitted models were Rational in the winter and Heat capacity in the summer.

#### 4.3.4 Growth rate

Growth rates were also highly variable among individuals at all temperatures, resulting again in low power of analysis (Fig. 7A). The optimum juvenile growth rate was measured between 21-27°C with maximum growth at 24°C (Fig. 7A). The ANOVA test revealed a significant difference between groups (P < 0.05) but the Bonferroni post-hoc test did not find significant differences between temperatures (Supplementary Table S15).

**Figure 7.**
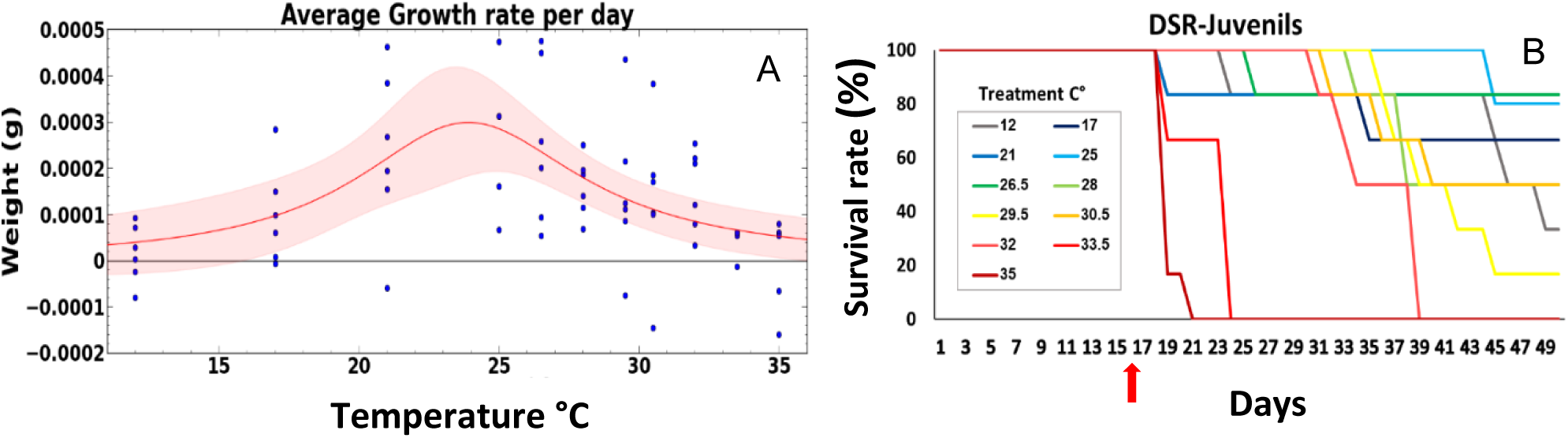
The effect of temperature on the growth rate. (A) **and on the daily survival rate** (DSR) (B) **in juvenile *P. elegans*** measured in the spring at 11 temperature levels between 12-35°C (n=6 per temperature level). The red arrow represents the time of reaching the target temperatures. The best fitted model in A was the Rational model.

#### 4.3.5 Critical Temperature Maximum and warming tolerance

The CTMax was lower in winter (36.46°C) compared to summer (38.1°C, t-test Mann-Whitney Test, U = .000, Z = -3.341, P < 0.001, Supplementary Table S16). When considering MHT of 37.8°C (measured in pool 1 in July 2017), the warming tolerance for tidepools is only 0.3°C, but in the open coastal waters, it is much higher: 5.9°C.

### 4.4 Survivorship

In all three experiments, DSR differed significantly among temperatures (Kaplan-Meier, P < 0.001, Supplementary Tables S17-S19). *P. elegans* did not survive in the highest target temperatures. In the winter experiment, all shrimp died at 34°C within 4 days, and at 33°C within 3 days (Fig. 8A). At 32°C, 80% of the shrimp died by the end of the experiment. All other treatments showed survival > 50%. In the summer experiment (Fig. 8B) all the shrimp died at 35°C within 6 days and at 34°C within 9 days. At 32°C, 50% of the shrimp were dead at the end of the experiment. All other treatments showed survival > 50%. In the juvenile growth spring experiment, DSR was much lower since the experiment lasted two months (Fig. 7B). All juveniles died within 6 days at 35°C, within 11 days at 33.5°C and within 27 days at 32°C. At 29.5°C, 80% of the shrimp died by the end of the experiment. At 12°C, 60% of the shrimp died by the end of the experiment and at 30.5°C and 28°C, 50% survived until the end of the experiment. All other treatments showed survival rates > 50%.

**Figure 8.**
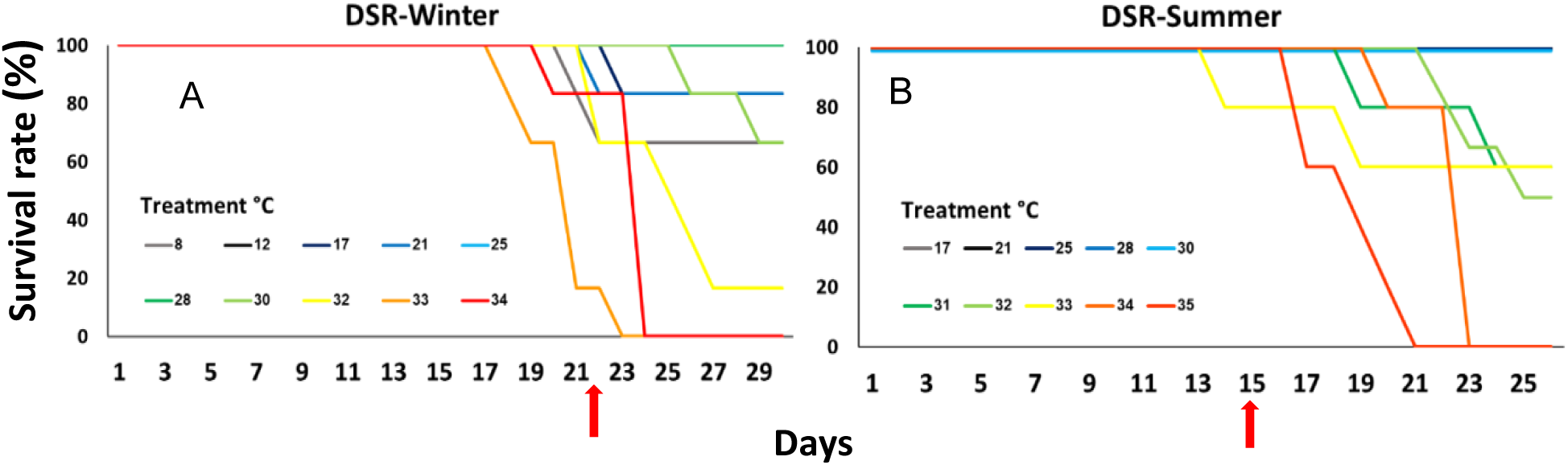
*P. elegans* daily survival rate (DSR) under 10 tested temperatures in the winter (**A**) and summer (**B**) experiments (n=6 per temperature level). The red arrows represents the time it took to reach the target temperatures. Each color represents a different treatment.

## 5. Discussion

Using multiple lines of evidence our results show that *Palaemon elegans* lives close to its upper thermal limits in the most thermally-stressful tidepools (small and often disconnected from the open sea), but it is still within a wide safe zone in more open habitats. Nonetheless, long exposures to temperatures slightly higher than current summer values may affect its population viability over time. We also found that the shrimp’s CTMax in the southeast Mediterranean, the warmest part of its distribution range, is higher compared to that of populations from the colder Atlantic shores, and that summer populations have higher thermal tolerance than winter ones, confirming our two working hypotheses related to local adaptation and seasonal acclimatization.

### 5.1 Tidepool conditions and population dynamics

As expected, our tidepool temperature time series revealed a wide daily variation of 10°C and seasonal variation of more than 32°C in the smaller, more isolated pools. Water temperature in these pools can drop to 6°C (after a few days of cold eastern winds) or climb up to 37.8°C and stay at temperatures greater than 35°C for more than six hours in summer. A few meters away, the open coastal waters have much lower temperature variability, showing a daily variation of 2-3°C, and seasonal variations of 17°C. In July and August, the open water temperature stayed above 30°C for more than two months, indicating the potential stress of residing in a shallow water system. Nonetheless, *P. elegans* populations were present in the tidepools throughout the year at variable densities and were more abundant at the higher part of the intertidal zone where the tidepools are less connected to the open sea and have the highest temperature fluctuations. The highest density of shrimp occurred during the recruitment season in the spring in the most isolated tidepools, similar to the findings of Vinagre et al (2015) who suggested that these tidepools may function as safe nursery grounds.

### 5.2 Respiration response to warming

The respiration rate of the shrimp increases with temperature until it levels off (or even slightly declines under long exposure in the summer), and when dead individuals are included in the analysis, clear optimum and lethal temperatures are evident (Fig. 4). As we hypothesized, respiration rates increase (and P_max_) from winter to summer, and T_opt_ also increases between seasons (28.2°C to 32°C), and so does CTMax (34.5°C to 35.5°C), suggesting seasonal acclimatization for the higher summer temperatures. Seasonal acclimatization of oxygen consumption rates has been previously documented in intertidal crabs (Newell, 1973). Hopkin et al (Hopkin et al., 2006) tested eight crabs species and in most, summer-captured animals had significantly higher CTMax values than winter captured animals. Evidence of seasonal acclimatization in cardiac responses to heat stress and thermal tolerance are commonly found in the literature. For instance, the temperature for peak heart rate in the gastropod, *Lymnaea stagnalis* (Harrison, 1977) was found to be lower in winter (30.5°C) than in summer (37.5°C). A recent study found a clear seasonal acclimatization of the cardiac function of the limpet *Patella vulgata*, that allows individuals to be more tolerant to higher temperatures in the warm months (Chapperon et al., 2016). A fast seasonal physiological acclimatization is a valuable feature that may lead to adaptive shifts in thermal breadth, thermal optima and higher fitness (Somero, 2002). Acclimatization is a known mechanism that enables the organism to cope with the changing environmental temperature that occurs seasonally. Acclimatization (in the wild) or acclimation (in the laboratory) can change the organism’s response to environmental conditions (Fangue et al., 2009; Lachenicht et al., 2010; Ravaux et al., 2016; Somero, 2002; Tomanek, 2010). For *P. elegans*, this acclimatization is critical, as summer shallow waters have already warmed considerably, by 2-3 degrees, in the past three decades (Ozer et al., 2016; Rilov, 2016). The duration (and frequency) of exposure to extreme conditions is also a critical parameter that can determine a species’ survival or fitness. In both winter and summer, *P. elegans* showed a reduction in the TPC parameters including P_max_ and T_opt_, with increased exposure to high temperatures, and thus support our third working hypothesis. *P. elegans* use energy to produce stress proteins for repairing temperature-induced damage (Madeira et al., 2015; Madeira et al., 2012a; Williams et al., 2016). Therefore, the long exposure to constant high temperatures probably has high demands on the shrimp’s metabolic functions and the accumulated stress weakens it, reducing its performance and shifting its thermal breadth to the left, as evident in our study. This is also expressed in the increased mortality at the highest temperatures after extended exposure.

The summer T_opt_ represents a threshold for respiration at 32°C. Beyond T_opt_, the rate drops sharply to zero. At 35°C mortality was 100% in six days and at 34°C, all shrimp died before the third incubation (less than ten days). In the wild, in tidepools, *P. elegans* experience much higher temperatures, above 35°C, but only for short periods of 6-7 hours maximum (with cooling at night, Fig. 2). Animals living in the intertidal zone develop different mechanisms to cope with short-term extreme temperatures. Some animals modify their exposure to extremes behaviorally, through the selection of thermally favorable microclimates, by hiding in crevices and escaping to shady areas (behavioral thermoregulation). Others have physiological adaptations where they divert their energy flow towards the production of a cellular stress response (CSR) and shift to anaerobic metabolism that is less efficient in generating energy (Han et al., 2013; Somero, 2002). These mechanisms require a change in the energy budget, which comes at the expense of growth, reproduction and other vital functions (Williams et al., 2016). With an expected 2 degrees increase in temperature in the future, and the subsequent increase in the duration and frequency of hot days which are expected to rise, we predict that in isolated tidepools CTMax will be frequently surpassed, leading to increased *P. elegans* mortality.

In the open coastal waters, at summer time, *P. elegans* currently experience temperatures that are constantly above 30°C for more than two months (average August temperatures around 31.5°C). Based on our results, an increase of two degrees in sea surface temperature (SST) in the open sea will lead to a severe reduction in individual performance and fitness and possibly in the population’s viability in the region. A total collapse of a regional populations due to warming (and possibly competition for food with an invasive species) has indeed already occurred in southeast Levant urchins (Yeruham et al., 2015; Yeruham et al., 2019). It is possible that semi-isolated tidepools might still offer temporal thermal refuge at night that can sustain the resilience of individuals living there and thus the population in the region. In future experiments, a simulation of real tidepool fluctuating conditions will shed more light on the actual robustness of the shrimp. Nonetheless, we assume that in tidepools disconnected from the open sea for long periods, multiple exposures to extreme peak temperatures in the heat of the day can gradually erode resilience – another aspect that deserves testing. The highest mortality that we witnessed in the field actually followed a sharp drop in temperature (down to 6°C); thus, cold stress in insulated pools during cold nights might also be a factor.

### 5.3 Gut resident time and activity respond to warming

Gut resident time reduced with temperature and was higher in winter than summer for the same temperatures, suggesting again a seasonal acclimatization in metabolic functions (faster digestion). Another crustacean study showed that at higher temperatures, there is a disassociation between feeding and gut evacuation rates and suggested that mechanisms other than digestion rate might limit feeding rate (Chipps, 1998). In our study, we did not test the feeding rates of *P. elegans*, but we did observe a decrease in appetite at high temperatures (personal observations), even though we found high rates of digestion at such temperatures in both seasons. These results suggest that an increased metabolic rate at higher water temperatures plus low feeding rates and high gut evacuation rates at the highest livable temperatures, likely reduce the overall scope for growth under excessive warming.

The activity test suggests again that higher metabolic rates, in summer compared to winter, are manifested with increased movement. There was a rise in the performance with temperature, but individual variability was very high, resulting in non-significant relationships. The higher metabolic rates in summer may express seasonal acclimatization to high temperatures.

### 5.4 Critical temperature maximum

The CTMax is considered to be the most reliable parameter in macrophysiological comparative studies in ectotherms (Lutterschmidt and Hutchison, 1997). It is used for assessing the upper thermal limits and the thermal tolerance of a species in relation to the maximum habitat temperature that a species experience. The CTMax, the thermal tolerance and the acclimation capacity for *P. elegans* were previously tested on the other side of the species distribution range, in the West Mediterranean and East Atlantic-which are colder regions compared to the EMS (Table 1). Madeira et al. (2012b) measured the CTMax of coastal and estuarine organisms including *P. elegans* on the coast of Portugal, where the mean water temperature was 24°C and MHT was 35°C, and established a CTMax of 34°C. The authors concluded that “the most vulnerable organisms to sea warming were those that occur in the thermally unstable environments because, despite their high CTMax values, they live closer to their thermal limits and have limited acclimation plasticity". These results were based on MHT measured in air and therefore their conclusion should be considered with caution, as it was not based on the actual MHT that *P. elegans* experience (in tidepool waters, which might be lower than that of the air). Vinagre et al. (2016) measured the CTMax and acclimation capacity of tropical and temperate coastal organisms, among them *P. elegans* from west Portugal. They found a CTMax of 33.4°C after 7 day exposure at ambient temperature (20°C). When testing the acclimation capacity of its thermal limits after acute and chronic exposure, they found that *P. elegans* has the highest acclimation capacity under long-term low stress (30 days +3°C, CTMax = 36.2°C) or short-term high stress (10 days +6°C, CTMax = 36.6°C) conditions compared to all other test species. Because in their study the MHT measured in a tidepool on a hot summer day was 30°C, they concluded that *P. elegans* is still far from its upper thermal limits in its natural environment.

In the western Mediterranean, Ravaux et al. (2016) examined the CTMax of *P. elegans* during ontogeny, and found that adults have the ability to adjust their CTMax through long-term acclimation (4 to 8 months at 10°C or 20°C). Although they used a slightly different procedure (heating rate of 1°C per minute) they found that the CTMax for zoea larva is significantly lower than that of adults, both reared at 20°C. The CTMax for adults acclimated to 10°C and 20°C was 27±0.8°C and 33.2±1.5°C, respectively. The higher value of CTMax for adult *P. elegans* is very similar to the CTMax of *P. elegans* measured in the study by Vinagre et al. (2016). Together, these findings indicate that the CTMax is strongly influenced by habitat temperatures experienced by the species.

Evidently, the CTMax values measured in our study for *P. elegans* at its trailing, warm edge (winter, 36.4°C; summer, 38.1°C) was considerably higher than those measured in populations from the leading edge of its distribution, as described above, even after acute exposure to extreme heat shown by Vinagre et al. (2016). The higher summer CTMax value (compared to winter) yet again is probably the result of acclimatization to the very high temperatures that this species experiences during this season. This result supports our hypothesis that the CTMax of *P. elegans* from this warmer region will be higher compared to that of western populations. It also supports the assumption that the upper thermal limit is related to the local thermal habitat. The mechanism behind it may be adaptation (perhaps through selection) to the higher temperatures in the region. Local adaptation was shown to be widespread in marine invertebrates including ones with planktonic larvae (Sanford and Kelly, 2011), such as *Palaemon*. However, isolated populations with high local adaptation might have very limited capacity to adapt to further warming, as was shown at the leading edge of a copepod that lives in high intertidal tidepools (Kelly et al., 2012).

### Juvenile growth

The optimum temperature for juvenile growth in the EMS was found to be 24°C (similar to late spring SST in coastal waters), and growth rates were highest between 21-27°C. In the field, juveniles were abundant in May and June, and the average temperature in pool 1, that was populated by the highest abundance of juveniles, during that period was 25.7°C and 29.3°C respectively, indicating that May has the average optimal temperature for growth. During that period, the maximum temperature recorded was 33.2°C (May) and 36.3°C (June); however, these extreme temperatures were short in duration and thus did not result in mass juvenile mortality. The survival data (Fig. 7B) showed that juvenile shrimp are more vulnerable to high and low temperatures compared to adults. Post-larva of *P. elegans* from the northwest Mediterranean were found to have lower CTMax than adults, based on their capacity for heat shock response (see, Ravaux et al., 2016). In our study, all juveniles died above 33°C, and mortality was greater than 50% at temperatures of 12, 28, 29.5 and 30°C.

In summary, *P. elegans* appears to have a high capacity for acclimatization (high plasticity) both biogeographically (although selection for more heat resistant genotypes might be involved in the process), and seasonally. *P. elegans* in the EMS has higher CTMax than its more northwestern counterparts, but in small, isolated tidepools it experiences short exposures to extreme temperatures that are much closer to the CTMax than those in the western populations. In light of the projected continuation of global warming of both air and water (IPCC, 2014), conditions in temporarily isolated tidepools are expected to become more extreme for *P. elegans* and especially for juveniles, which may negatively affect the population along the coast. Furthermore, the recently described prolonged desiccation events (PDEs) on southeastern Mediterranean rocky shores, generated by dry easterly winds that can expose the vermetid reefs to air continuously for days during spring (Zamir et al., 2018), can leave many tidepools isolated from the open sea for very long periods, creating extreme conditions in them that are harmful for juvenile shrimp. The frequency of PDE generating synoptic systems has more than doubled in the past four decades during winter and spring (Alpert et al., 2004; Zamir et al., 2018), and this trend is expected to continue in the near future (Hochman et al., 2018a; Hochman et al., 2018b), adding additional stress to this already challenging habitat. The combination of all these stressors has the potential to reduce the fitness and viability of the population on a regional scale with potentially little chance of adaptation as warm-tolerant phenotypes in the trailing edge could reduce genetic diversity and constrain the potential for adaptation for other stressors (Donelson et al., 2019).

The potential for plasticity in thermal tolerance, and/or adaptation, which may mitigate the effects of global warming is largely unknown due to the lack of knowledge about global and taxonomic patterns of variation in tolerance plasticity (Gunderson and Stillman, 2015). Donelson et al (2019) further stress the importance of such knowledge to improve species distribution models under climate change and for marine conservation. Gunderson and Stillman’s (2015) global analysis of terrestrial, freshwater and marine ectotherms, suggested that “overheating risk will be minimally reduced by acclimation in even the most plastic groups”. The findings of our study strengthen the case for the clear context-dependency of the vulnerability of ectothermic species that live in extremely stressful and fluctuating environments to climate change. It demonstrates the potential biogeographical thermal physiological flexibility and adaptability (whether genotypic or phenotypic) of such species, but also their high and increasing vulnerability at the trailing edge of their population under climate change as suggested by Somero (2010).

## Acknowledgments

We would like to thank for the hard work of the Rilov Lab technicians, D. Golomb, O. Peleg, and M. Schonwald, as well as many students including T. Guy-Haim, E. Yeruham, M. Mulas, and two fellow researches, J. Silverman and E. Rahav. The work was supported by the Israeli Science Foundation (ISF) grant number 117/10 (to GR), and the by the Ministry of Science & Technology of the State of Israel and Federal Ministry of Education and Research (BMBF), Germany, grant number 9732 (to GR).

## References

Al-Khafaji, K. K., Al Qarooni, I. H., Al Abbad, M. Y. and Al-Lateef, N. M. A. (2016). Study of the growth, reproductive biology and abundance for invasive shrimps Palaemon elegans Rathke from Garmat Ali river Basrah, Southern Iraq. Journal of Coastal Life Medicine 4, 536-540.10.12980/jclm.4.2016J6-97

Alpert, P., Osetinsky, I., Ziv, B. and Shafir, H. (2004). A new seasons definition based on classified daily synoptic systems: an example for the eastern Mediterranean. International Journal of Climatology: A Journal of the Royal Meteorological Society 24, 1013-1021.10.1002/joc.1037

Angilletta Jr, M. J. (2006). Estimating and comparing thermal performance curves. Journal of Thermal Biology 31, 541–545

Bates, A. E., Pecl, G. T., Frusher, F., Hobday, A. J., Wernberg, T., Smale, D. A., Sunday, J. M., Hill, N. A., Dulvy, N. K. and Colwell, K. (2014). Defining and observing stages of climate-mediated range shifts in marine systems. Global Environmental Change 26, 27–38

Bilgin, S., Ozen, O. and Samsun, O. (2009). Sexual seasonal growth variation and reproduction biology of the rock pool prawn, Palaemon elegans (Decapoda: Palaemonidae) in the southern Black Sea. Scientia Marina 73, 239–247

Chapman, B. B., Hegg, A. and Ljungberg, P. (2013). Sex and the syndrome: individual and population consistency in behaviour in rock pool prawn Palaemon elegans. PLoS One 8

Chapperon, C., Volkenborn, N., Clavier, J., Séité, S., Seabra, R. and Lima, F. P. (2016). Exposure to solar radiation drives organismal vulnerability to climate: evidence from an intertidal limpet. Journal of Thermal Biology 57, 92–100

Chipps, S. R. (1998). Temperature-dependent consumption and gut-residence time in the opossum shrimp Mysis relicta. Journal of plankton research 20, 2401–2411

Dalla Via, J. (1985). Oxygen consumption and temperature change in the shrimp Palaemon elegans. Marine Ecology Progress Series 26, 199–202

DeMaster, D. P. and Drevenak, J. K. (1988). Survivorship patterns in three species of captive cetaceans. Marine Mammal Science 4, 297–311

Donelson, J. M., Sunday, J. M., Figueira, W. F., Gaitán-Espitia, J. D., Hobday, A. J., Johnson, C. R., Leis, J. M., Ling, S. D., Marshall, D. and Pandolfi, J. M. (2019). Understanding interactions between plasticity, adaptation and range shifts in response to marine environmental change. Philosophical Transactions of the Royal Society B 374, 20180186

Duran, M., Suicmez, M., Kayim, M. and Kaynar, C. (2006). Preliminary analysis of the biological characteristics of Palaemon elegans (Decapoda, Palaemonidae) in the Coast of Sinop, Black Sea, N. Turkey. Pakistan Journal of Biological Sciences 9, 848–853

Fangue, N. A., Richards, J. G. and Schulte, P. M. (2009). Do mitochondrial properties explain intraspecific variation in thermal tolerance? Journal of Experimental Biology 212, 514–522

Fofonoff, P., Ruiz, G., Steves, B. and Carlton, J. (2003). National exotic marine and estuarine species information system. Web publication< http://invasions.si.edu/nemesis7380

Gattuso, J.-P., Magnan, A., Billé, R., Cheung, W., Howes, E., Joos, F., Allemand, D., Bopp, L., Cooley, S. and Eakin, C. (2015). Contrasting futures for ocean and society from different anthropogenic CO^2^ emissions scenarios. Science 349, 4722-1-4722-10.10.1126/science.aac4722

Glamuzina, L., Conides, A., Prusina, I., Ćukteraš, M., Klaoudatos, D., Zacharaki, P. and Glamuzina, B. (2014). Population structure, growth, mortality and fecundity of Palaemon adspersus (Rathke 1837; Decapoda: Palaemonidae) in the Parila Lagoon (Croatia, SE Adriatic Sea) with notes on the population management. Turkish Journal of Fisheries and Aquatic Sciences 14, 677–687

Gunderson, A. R. and Stillman, J. H. (2015). Plasticity in thermal tolerance has limited potential to buffer ectotherms from global warming. Proceedings of the Royal Society B: Biological Sciences 282, 20150401

Han, G.-d., Zhang, S., Marshall, D. J., Ke, C.-h. and Dong, Y.-w. (2013). Metabolic energy sensors (AMPK and SIRT1), protein carbonylation and cardiac failure as biomarkers of thermal stress in an intertidal limpet: linking energetic allocation with environmental temperature during aerial emersion. Journal of Experimental Biology 216, 3273–3282

Harley, C. D. G., Hughes, A. R., Hultgren, K. M., Miner, B. G., Sorte, C. J. B., Thornber, C. S., Rodriguez, L. F., Tomanek, L. and Williams, S. L. (2006). The impacts of climate change in coastal marine systems. Ecology letters 9, 228–241

Harrison, P. (1977). Seasonal changes in the heart rate of the freshwater pulmonate Lymnaea stagnalis (L.). Comparative Biochemistry and Physiology Part A: Physiology 58, 37–41

Helmuth, B., Mieszkowska, N., Moore, P. and Hawkins, S. J. (2006). Living on the edge of two changing worlds: Forecasting the responses of rocky intertidal ecosystems to climate change. Annual Review of Ecology Evolution and Systematics 37, 373-404.10.1146/annurev.ecolsys.37.091305.110149

Hochman, A., Harpaz, T., Saaroni, H. and Alpert, P. (2018a). The seasons’ length in 21st century CMIP5 projections over the eastern Mediterranean. International Journal of Climatology 38, 2627–2637

Hochman, A., Mercogliano, P., Alpert, P., Saaroni, H. and Bucchignani, E. (2018b). High-resolution projection of climate change and extremity over Israel using COSMO-CLM. International Journal of Climatology 38, 5095–5106

Hopkin, R. S., Qari, S., Bowler, K., Hyde, D. and Cuculescu, M. (2006). Seasonal thermal tolerance in marine Crustacea. Journal of Experimental Marine Biology and Ecology 331, 74–81

IPCC. (2014). Climate Change 2014: Impacts, Adaptation, and Vulnerability. Part A: Global and Sectoral Aspects. Contribution of Working Group II to the Fifth Assessment Report of the Intergovernmental Panel on Climate Change, 6914 eds. R. K. Pachauri and L. A. Meyer), pp. 151 pp. Geneva, Switzerland: IPCC.

IPCC. (2019). Summary for Policymakers. In: IPCC Special Report on the Ocean and Cryosphere in a Changing Climate, 7325 eds. H. O. Pörtner D. C. Roberts V. Masson-Delmotte P. Zhai M. Tignor E. Poloczanska K. Mintenbeck M. Nicolai A. Okem J. Petzold et al.).

Janas, U. and Mańkucka, A. (2010). Body size and reproductive traits of Palaemon elegans Rathke, 1837 (Crustacea, Decapoda), a recent colonizer of the Baltic Sea. Oceanological and Hydrobiological Studies 39, 3–24

Janas, U., Piłka, M. and Lipińska, D. (2013). Temperature and salinity requirements of Palaemon adspersus Rathke, 1837 and Palaemon elegans Rathke, 1837. Do they explain the occurrence and expansion of prawns in the Baltic Sea? Marine Biology Research 9, 293–300

Kelly, M. W., Sanford, E. and Grosberg, R. K. (2012). Limited potential for adaptation to climate change in a broadly distributed marine crustacean. Proceedings of the Royal Society B: Biological Sciences 279, 349–356

Lachenicht, M., Clusella-Trullas, S., Boardman, L., Le Roux, C. and Terblanche, J. (2010). Effects of acclimation temperature on thermal tolerance, locomotion performance and respiratory metabolism in Acheta domesticus L.(Orthoptera: Gryllidae). Journal of Insect Physiology 56, 822–830

Levinton, J. (2001). The Chemical and Physical Environment. In Marine Biology, 7390, pp. 592: Oxford University Press.

Lima, F. P., Gomes, F., Seabra, R., Wethey, D. S., Seabra, M. I., Cruz, T., Santos, A. M. and Hilbish, T. J. (2016). Loss of thermal refugia near equatorial range limits. Global change biology 22, 254–263

Lutterschmidt, W. I. and Hutchison, V. H. (1997). The critical thermal maximum: history and critique. Canadian Journal of Zoology 75, 1561–1574

Macias, D., Garcia-Gorriz, E. and Stips, A. (2013). Understanding the causes of recent warming of Mediterranean waters. How much could be attributed to climate change? PLoS One 8, e81591

Madeira, D., Mendonça, V., Dias, M., Roma, J., Costa, P. M., Larguinho, M., Vinagre, C. and Diniz, M. S. (2015). Physiological, cellular and biochemical thermal stress response of intertidal shrimps with different vertical distributions: Palaemon elegans and Palaemon serratus. Comparative Biochemistry and Physiology Part A: Molecular & Integrative Physiology 183, 107–115

Madeira, D., Narciso, L., Cabral, H., Vinagre, C. and Diniz, M. (2012a). HSP70 production patterns in coastal and estuarine organisms facing increasing temperatures. Journal of Sea Research 73, 137–147

Madeira, D., Narciso, L., Cabral, H. N. and Vinagre, C. (2012b). Thermal tolerance and potential impacts of climate change on coastal and estuarine organisms. Journal of Sea Research 70, 32–41

Magozzi, S. and Calosi, P. (2015). Integrating metabolic performance, thermal tolerance, and plasticity enables for more accurate predictions on species vulnerability to acute and chronic effects of global warming. Global change biology 21, 181–194

Marbà, N., Jorda, G., Agusti, S., Girard, C. and Duarte, C. M. (2015). Footprints of climate change on Mediterranean Sea biota. Frontiers in Marine Science 2.10.3389/fmars.2015.00056

Mora, C. and Ospina, A. (2001). Tolerance to high temperatures and potential impact of sea warming on reef fishes of Gorgona Island (tropical eastern Pacific). Marine Biology 139, 765–769

Newell, R. C. (1973). Factors affecting the respiration of intertidal invertebrates. American zoologist 13, 513–528

Ozer, T., Gertman, I., Kress, N., Silverman, J. and Herut, B. (2016). Interannual thermohaline (1979–2014) and nutrient (2002–2014) dynamics in the Levantine surface and intermediate water masses, SE Mediterranean Sea. Global and Planetary Change 6281 10.1016/j.gloplacha.2016.04.001.10.1016/j.gloplacha.2016.04.001

Pastor, F., Valiente, J. A. and Palau, J. L. (2019). Sea surface temperature in the Mediterranean: Trends and spatial patterns (1982–2016). In Meteorology and Climatology of the Mediterranean and Black Seas, vol. Pageoph Topical Volumes eds. Vilibić K. Horvath and J. Palau), pp. 297–309. Cham: Birkhäuser.

Pörtner, H. O. and Farrell, A. P. (2008). Physiology and climate change. Science 322, 690–692

Ravaux, J., Léger, N., Rabet, N., Fourgous, C., Voland, G., Zbinden, M. and Shillito, B. (2016). Plasticity and acquisition of the thermal tolerance (upper thermal limit and heat shock response) in the intertidal species Palaemon elegans. Journal of Experimental Marine Biology and Ecology 484, 39–45

Rilov, G. (2016). Multi-species collapses at the warm edge of a warming sea. Scientific reports 6, 36897. 10.1038/srep36897

Rivetti, I., Fraschetti, S., Lionello, P., Zambianchi, E. and Boero, F. (2014). Global warming and mass mortalities of benthic invertebrates in the Mediterranean Sea. PLoS One 9. 10.1371/journal.pone.0115655

Sanford, E. and Kelly, M. W. (2011). Local Adaptation in Marine Invertebrates. In Annual Review of Marine Science, Vol 3, vol. 3, pp. 509–535.

Schulte, P. M., Healy, T. M. and Fangue, N. A. (2011). Thermal performance curves, phenotypic plasticity, and the time scales of temperature exposure. Integrative and comparative biology 51, 691–702

Seebacher, F., White, C. R. and Franklin, C. E. (2015). Physiological plasticity increases resilience of ectothermic animals to climate change. Nature Climate Change 5, 61–66

Somero, G. (2010). The physiology of climate change: how potentials for acclimatization and genetic adaptation will determine ‘winners’ and ‘losers’. Journal of Experimental Biology 213, 912–920

Somero, G. N. (2002). Thermal physiology and vertical zonation of intertidal animals: optima, limits, and costs of living. Integrative and comparative biology 42, 780–789

Szaniawska, A. and Lapinska, E. (2006). Environmental preferences of Crangon crangon (Linnaeus, 1758), Palaemon adspersus Rathke, 1837, and Palaemon elegans Rathke, 1837 in the littoral zone of the Gulf of Gdansk. Crustaceana 79, 649–662

Tomanek, L. (2010). Variation in the heat shock response and its implication for predicting the effect of global climate change on species’ biogeographical distribution ranges and metabolic costs. Journal of Experimental Biology 213, 971–979

Tsurnamal, M. (1963). Larval development of the prawn Palaemon elegans Rathke (Crustacea, Decapoda) from the coast of Israel. Israel Journal of Zoology 12, 117–141

Verberk, W. C., Bartolini, F., Marshall, D. J., Pörtner, H. O., Terblanche, J. S., White, C. R. and Giomi, F. (2016). Can respiratory physiology predict thermal niches? Annals of the New York Academy of Sciences 1365, 73–88

Vinagre, C., Dias, M., Fonseca, C., Pinto, M. T., Cabral, H. N. and Silva, A. (2015). Use of rocky intertidal pools by shrimp species in a temperate area. Biologia 70, 372–379

Vinagre, C., Dias, M., Roma, J., Silva, A., Madeira, D. and Diniz, M. S. (2013). Critical thermal maxima of common rocky intertidal fish and shrimps—A preliminary assessment. Journal of Sea Research 81, 10–12

Vinagre, C., Leal, I., Mendonça, V., Madeira, D., Narciso, L., Diniz, M. S. and Flores, A. A. (2016). Vulnerability to climate warming and acclimation capacity of tropical and temperate coastal organisms. Ecological indicators 62, 317–327

Vinagre, C., Mendonca, V., Cereja, R., Abreu-Afonso, F., Dias, M., Mizrahi, D. and Flores, A. A. (2018). Ecological traps in shallow coastal waters—Potential effect of heat-waves in tropical and temperate organisms. PLoS One 13

Williams, C. M., Buckley, L. B., Sheldon, K. S., Vickers, M., Pörtner, H.-O., Dowd, W. W., Gunderson, A. R., Marshall, K. E. and Stillman, J. H. (2016). Biological impacts of thermal extremes: mechanisms and costs of functional responses matter. Integrative and comparative biology 56, 73–84

Yeruham, E., Rilov, G., Shpigel, M. and Abelson, A. (2015). Collapse of the echinoid *Paracentrotus lividus* populations in the Eastern Mediterranean - result of climate change? Scientific reports 5, 13479. 10.1038/srep13479

Yeruham, E., Shpigel, M., Abelson, A. and Rilov, G. (2019). Ocean warming and tropical invaders erode the performance of a key herbivore. Ecology 7280 10.1002/ecy.2925.10.1002/ecy.2925

Zamir, R., Alpert, P. and Rilov, G. (2018). Increase in weather patterns generating extreme desiccation events: implications for Mediterranean rocky shore ecosystems. Estuaries and Coasts 41, 1868–1884.10.1007/s12237-018-0408-5

